# Structural connectivity-informed dynamic estimation (STRiDE): Multimodal connectivity constrained ICA

**DOI:** 10.1101/2025.06.18.660402

**Authors:** Mahshid Fouladivanda, Armin Iraji, Lei Wu, Jiayu Chen, Pablo Andrés Camazón, Theodorus G.M. van Erp, Aysenil Belger, Godfrey D. Pearlson, Tulay Adali, Vince D. Calhoun

**Affiliations:** Tri-institute Translational Research in Neuroimaging and Data Science (TReNDS Center), Georgia State University, Georgia Institute of Technology, Emory University, Atlanta, GA, USA; Georgia State University, Atlanta, GA, USA; Institute of Psychiatry and Mental Health Gregorio Marañón, Madrid, Spain; Clinical Translational Neuroscience Laboratory, Department of Psychiatry and Human Behavior School of Medicine, University of California, Irvine, CA, USA; Department of Psychiatry Director, Neuroimaging Research in Psychiatry Director, Clinical Translational Core, UNC Intellectual and Developmental Disabilities Research Center, University of North Carolina, Chapel Hill, NC, USA; Olin Neuropsychiatry Research Center, Department of Psychiatry and Neuroscience, Yale University, School of Medicine, New Heaven, CT, USA; Department of CSEE, University of Maryland Baltimore County, Baltimore, MD, USA

**Keywords:** Structural connectivity, Dynamic functional connectivity, Connectivity-constrained ICA, Multimodal brain states, Optimization

## Abstract

Understanding the dynamic intrinsic interactions between brain regions has been advanced by functional magnetic resonance imaging (fMRI), particularly through the connectivity analysis used to characterize reoccurring patterns in the brain known as brain states. However, previous studies have primarily focused on unimodal models, which can hinder optimal dynamic state estimation, especially in cognitive disorders, cases where there may be both structure and function disruption. To better estimate intrinsic brain interactions, it is important to account for the factors shaping this estimation not only in terms of time resolved variability of the connectivity but also regarding the underlying physical pathways between brain regions. However current approaches mostly use deterministic weighting of structural connectivity. To address this, we propose a flexible multimodal connectivity constrained independent component analysis (ICA) model, termed structural connectivity-informed dynamic state estimation (STRiDE), that enhances stability and sensitivity by leveraging white matter structural connectivity and dynamic functional connectivity information. Using this model, we decompose brain interactions into independent, reoccurring multimodal patterns or structural-functional states, guided by maximally independent structural connectivity priors derived from the group level data. We first evaluate our proposed model using a simulation pipeline, showing the approach works as design and improves sensitivity to group differences and enhances robustness to noise. Next, we applied the proposed multimodal model to real dataset including a cohort of subjects with schizophrenia (SZ) and healthy controls (HC). Results demonstrated its potential to enhance group-differences in both connectivity domain and temporal dynamics parameters. Specifically results highlighted disruption within and between sensory and trans-modal domains, through a SZ vs HC comparison. Symptoms severity and cognitive scores statistical analysis specifies their significant association with default mode domain, offering insights into the disrupted functional and neural mechanisms underlying schizophrenia. In addition, temporal interplay of the estimated STRiDEs reveals that visual-related

STRiDE is significantly impacted in SZ, regarding the speed of processing score, underscoring the link between visual system and speed of processing. In sum, the STRiDE approach provides a flexible way to link structural and functional connectivity at the network level and represents a general approach for studying multimodal dynamic patterns and leveraging these to study the typical and disordered brain.

## Introduction

Resting-state functional magnetic resonance imaging (fMRI) has highlighted that brain activity exhibits dynamic characteristics during rest. Dynamic functional network connectivity (dFNC) analysis captures multiple recurring patterns of functional connectivity, [1, 2] known as brain states challenging the long-standing assumption of stationary in brain function [3-6]. As these brain states transition over time, they offer a richer understanding of cognitive and functional processes, particularly in individuals with cognitive impairments and disorders such as schizophrenia [3, 4]. While, dynamic functional network connectivity provides critical insights into intrinsic brain function and its temporal properties [2, 7], understanding how these dynamics are shaped by the underlying structural architecture of the brain has emerged as a key area of research [8-12].

The human brain’s structural connectivity, known as connectome [13] is defined by the physical pathways connecting brain functional networks, serves as the scaffold for neural activity. Structural connectivity is typically derived from diffusion magnetic resonance imaging (dMRI) using tractography analysis. While other imaging modalities such as PD, T1, or T2 can provide structural information (e.g., gray matter volume), dMRI uniquely captures the white matter pathways responsible for information transfer. Structural connectivity analysis provides insights into the patterns and constraints that shape functional interactions [14] however its independent role in guiding dynamic brain processes remains underexplored [14-16]. Bridging this gap requires a deeper understanding of how these static foundations influence the brain’s functional dynamic.

Structural-functional dynamic modeling plays a critical role in understanding how the brain’s anatomical features contribute to its functional dynamics. These models provide a framework for linking structural elements, such as white matter pathways, to the temporal evolution of neural activity. Multimodal structural-functional modeling offers valuable insights into cognitive processes, behavior, and neuropathology, particularly in disorders that involve both structural and functional disruptions, such as schizophrenia [10, 17].

Recent work has focused on integrating multiple imaging modalities, particularly rs-fMRI and dMRI. However, there remains a need for novel approaches that can address key challenges, including the high dimensionality of neuroimaging data, disparate spatial and temporal resolutions, and variability in image types, scales, and formats [18]. Some efforts [12, 17, 19] have attempted to overcome these challenges by leveraging network connectivity domain analyses. While multimodal models in the connectivity domain show promise, they often rely on simple correlation or distance metrics to link structural and functional connectivity. Consequently, they fall short of capturing the inherently asymmetric and complex relationship between brain structure and function [20]. Alternatively, computational models [21, 22] have been developed to simulate brain functional activity based on structural connectivity. These models, although successful in generating plausible simulations, are typically based on group-level data and simplified assumptions to balance complexity with tractability [21, 22]. This limits their ability to capture individual variability and the broader neurochemical intricacies of brain function. Data-driven structural-functional models, particularly those employing independent component analysis (ICA) [19, 23, 24] have shown promise in improving our understanding of brain connectivity changes associated with clinical disorders [23]. Other approaches, such as connectome harmonic decomposition [10, 19], aim to uncover multiscale structure-function relationships by analyzing underlying connectivity patterns. While these methods account for distributed structural influences, their focus on global spatial frequencies may limit sensitivity to localized brain dynamics [10, 19].

Despite the promise of computational and multimodal approaches, many models still assume functional connectivity to be static over time. Although a limited number of studies have demonstrated the influence of structural connectivity on functional dynamics, they often rely on indirect approximations and typically incorporate structural information only as weights in dynamic models [25, 26]. However, there is growing recognition of the importance of integrating structural and dynamic functional information. A more comprehensive framework is needed to fully capture the interplay between structure and time-varying brain function.

To overcome existing limitations, we propose a novel multimodal, data-driven model that jointly integrates structural and dynamic functional connectivity information. Integrating these modalities within the connectivity domain addresses the high dimensionality of neuroimaging data. The proposed Structural-Informed Dynamic State Estimation (STRiDE) model leverages multiple independent structural bases, used as priors, to inform dynamic brain analysis at the individual level. We applied group ICA model [27] to extract a set of structural bases, which then guide the estimation of maximally independent functional states from resting-state fMRI. This is achieved by grounding dynamic connectivity analysis in the estimated structural bases within a data-driven, constrained ICA framework [28, 29]. Identifying structural bases in this way is a novel method for revealing independent white matter connectivity patterns, providing a low-dimensional representation of structural connectivity across individuals. Because structural and functional connectivity are not directly or symmetrically related, using a set of structural bases helps overcome existing challenges in integrating different imaging modalities. Moreover, the ICA-based model simplifies the overall framework while enabling joint learning from information with different spatial and temporal resolutions. Most importantly, our approach aims to offer new insights into the relationship between the brain’s physical architecture and its functional dynamics—an area that has not been thoroughly explored. While our previous work focused on integrating structural and static functional information [23], the present study identifies six distinct multimodal brain states, capturing dynamic functional patterns. Furthermore, this approach marks a departure from traditional methods by emphasizing the independent role of structural bases as precursors to dynamic analysis. The study concludes by demonstrating the utility of this framework in elucidating brain function and advancing our understanding of neural processes, paving the way for future research in neuroscience and related fields.

## Method

### 1. Model definition

We propose a multimodal structural and dynamic functional network learning model that extends the ICA framework by incorporating structural connectivity bases as constraints on dynamic functional network connectivity states, (see Figure 1.a-c). Figure 1.a presents the overall pipeline of the proposed model. We first extracted the structural connectivity bases (Figure 1.b) as described in paragraph **Structural connectivity bases.** Dynamic functional network connectivity is computed as detailed in subsection **3. Real data**. These estimated structural and dynamic functional connectivity features were then integrated through the multimodal model, as illustrated in Figure 1.c.

**Figure 1.**
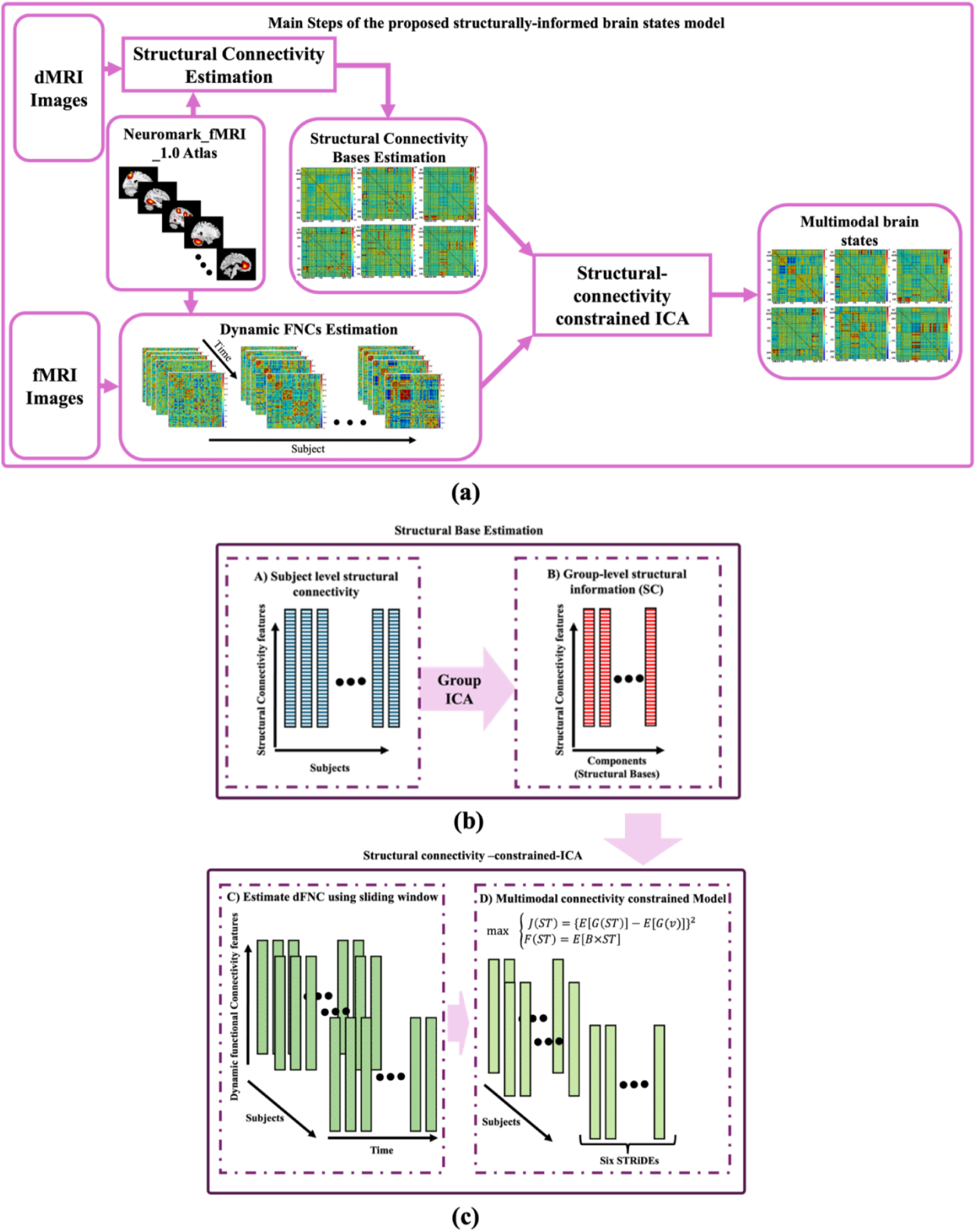
illustrated the pipeline of the proposed multimodal structural-informed dynamic states estimation model. In (a) the overall steps including from the inputs to outputs is shown. Preprocessed rs-fMRI and dMRI and NeuroMark_fMRI_1.0 atlas used as input. Using the NeuroMark_fMRI_1.0 atlas, subject-level structural connectivity and dynamic functional connectivity network (dFNC) were estimated. Next, we determined a set of structural bases using the group ICA (GICA) model, described in (b). finally, estimated structural bases along with the dFNCs were input to the structurally connectivity -constrained ICA model (as in (c)) to compute the multimodal brain states, resulting in the STRiDE model.

#### Structural connectivity bases

To estimate the structural connectivity bases, we first computed a *N* × *N* structural connectivity matrix for each subject (as described in subsection **3. Real data,** paragraph **Brain Network Connectivity**). These matrices were then flattened and concatenated over subjects (*M*). Next, we applied group ICA (GICA) [27, 30], as implemented in the GIFT toolbox (https://trendscenter.org/software/gift) [31]. GICA is a computational technique used to decompose a multivariate set of observations into additive components using different algorithms. Here, we used infomax algorithm, shown to yield reliable results [32]. It assumes that the observed data are linear mixtures of statistically independent source signals, and it aims to recover these sources from the observed mixtures, as described in Equation 1. Applying GICA to the structural connectivity data reveals independent structural connectivity patterns-referred to structural bases-that frequently occur across subjects. These bases capture consistent, coherent patterns of structural interconnectivity among brain functional networks. GICA also provides the weighted contribution of each structural base at different time for individuals (Figure 1.b)

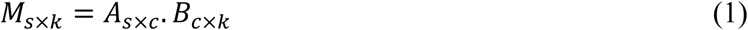

where, *M* represents the observed mixed data, i.e., the structural connectivity matrices, with dimensions corresponding to the number of subjects (*s*) by the number of the structural connectivity features (*k* = *N* × (*N* − 1)/2). B represents the independent sources, i.e., structural bases, while *A* is the mixing matrix. There are different ways to estimate the number of the independent sources (*c*). We used the elbow criteria implemented in GIFT to determine the number of structural bases, which suggested a model order of six.

#### Proposed Model

The proposed model is a multi-objective optimization process, leveraged from the constrained ICA model [28, 29]. In the constrained ICA model, a set of prior information, referred to as references, is used to guide the extraction of maximally independent components during the optimization process. Building on this concept, we developed a model to guide state estimation analysis from the dynamic functional connectivity (dFNCs) using the structural information as references. We termed this model structurally-informed dynamic estimation (STRiDE).

For each subject dFNCs were estimated as described in subsection **3. Real data**, paragraph **Brain Network Connectivity**

These dFNCs were then decomposed into independent patterns (STRiDE), which incorporate the structural references (structural bases) by optimizing the model through a dual-objective function defined in Equation 2.

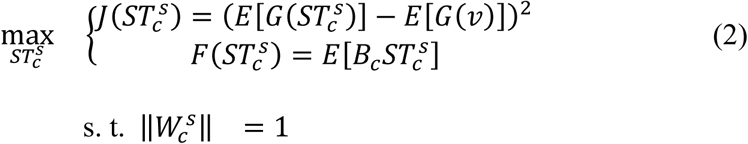

Here, *ST_c_^s^* denotes the estimated STRiDE for subject s, and c indexes the number of estimated STRiDEs. *E* is the expectation operator, *G* is any quadratic function, and *v* is a random variable. In Equation 2, *J*(.) can be negentropy or kurtosis [33] function, promoting statistical independence, while *F*(.) enforces similarity with the structural bases, acting as the constraint. To solve the multi-objective problem in Equation 2, we applied a common approach, a linear weighted sum of the objectives, simplifying the optimization as follows:

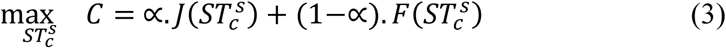

where, ∝ is a constant weight, set to 0.5. To prevent bias from differing objectives (cost functions), we normalize the cost functions in Equations 3 using the method described in {Du, 2013 #31}, and used equal weights. Then, the optimization was carried out using a gradient ascent method, iteratively converging to an optimal solution. Finally, we identified six STRiDEs for each subject and quantified their contributions over time—referred to as structurally-informed time courses.

#### Power and dynamic parameters of structurally informed brain states

In addition to the connectivity patterns (STRiDEs), a mixing matrix was estimated to identify the weighted contribution of each structural base across different dFNC matrices. This matrix was referred to as time courses (TC). To determine how STRiDEs evolve over time, we thresholded the TC by assigning the dominant STRiDE to a time interval. if its weight exceeded the median value of all weights.

In this study, we investigated the total power of the TC corresponding to each STRiDE. Total power is defined as the magnitude of each STRiDE’s contribution to the dFNCs regardless of sign, at each time interval, as shown in Equation 4.

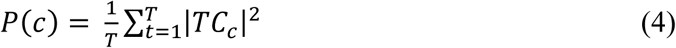

Here, T is the number of time intervals.

Furthermore, we examined the dynamic properties of brain states, characterized by four key temporal metrics including frequency, mean dwell time, occupancy rate and the transition probability matrix [1, 34]. Frequency refers to the number of times a subject transition into a given state throughout the scan. Mean dwell time captures the average duration (in time points) a subject remains continuously in a particular state before switching. Occupancy rate is defined as the proportion of total time spent in each state, reflecting the dominance of specific brain dynamics. Lastly, the transition matrix quantifies the probability of switching from one state to another, providing insights into the temporal organization and fluidity of state changes across the brain network.

### 2. Simulation

A hypothetical simulation was conducted to evaluate the proposed multimodal model by comparing the estimated STRiDEs with single-modal (functional) brain states. The underlying hypothesis is that incorporating structural information to guide state estimation enhances the accuracy of detected patterns in brain functional interactions. To test this hypothesis, we designed a simulation pipeline as described below.

First, two 50 × 50 matrices representing distinct brain sources—with different connectivity patterns across 50 regions or networks—were defined as ground truth (Figure 2.a). These two sources were then linearly combined using a mixing matrix to generate structural connectivity data for 10 individual subjects. The mixing matrix was randomly generated with a mean of zero and a standard deviation of one. One of the sources (source 1) was subsequently manipulated by randomly increasing or decreasing the weights of specific connections (target regions shown in Figure 2). Using these altered sources, structural connectivity for an additional 10 individuals was generated using the same procedure, simulating variability between two distinct groups.

**Figure 2.**
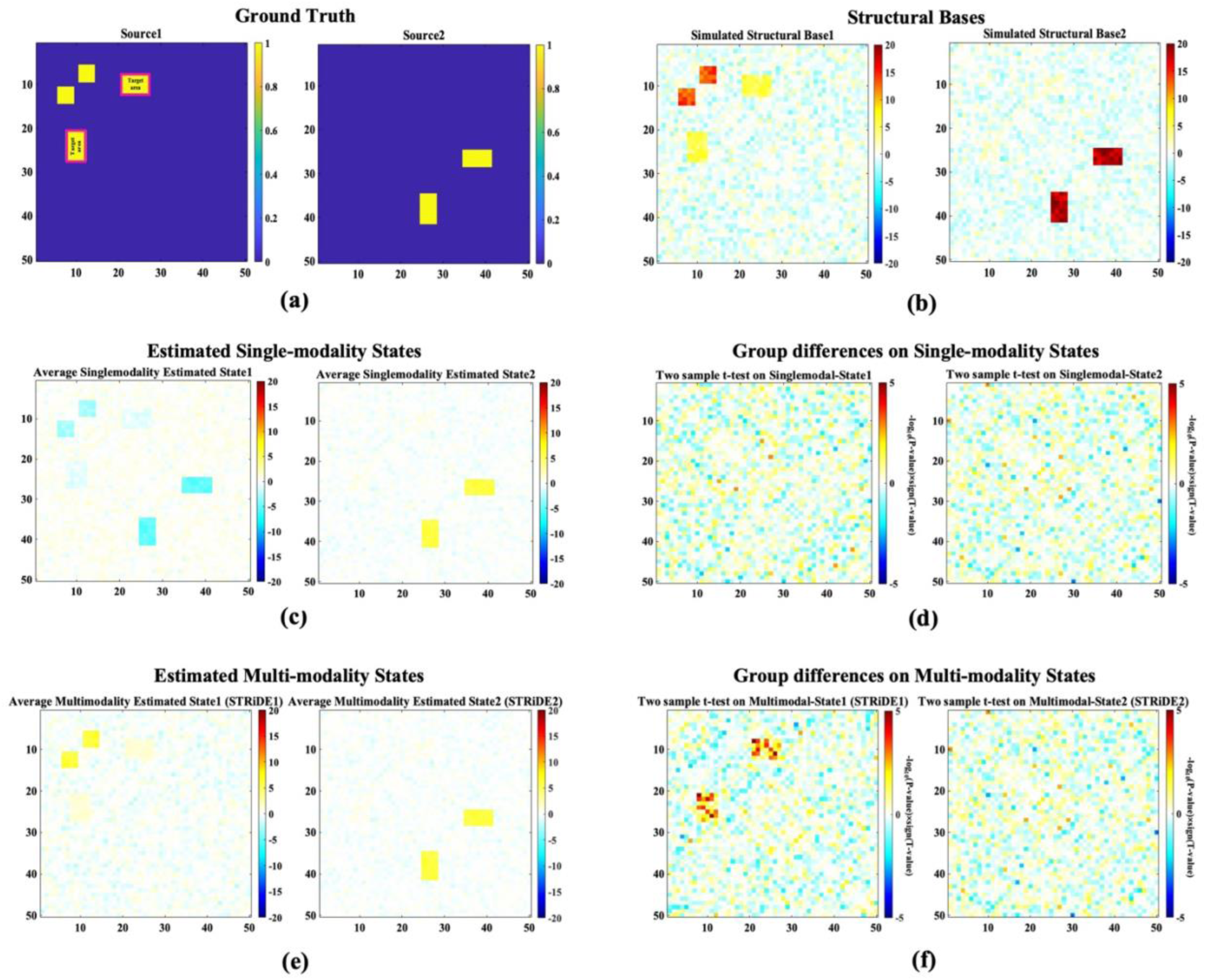
A simulation illustrating the STRiDE advantages. (a) A representative of two sources (connectivity pattern) used as ground truth. (b) Estimated simulated structural bases. (c) the average of the single-modal states estimated using only functional data. (d) two sample t-test on each single-modal state separately. Single-modal state1 and state2 show not significant connections between the group. (e) the average of the multi-modal states (STRiDE) estimated using both structural and functional data. (f) two sample t-test on each multi-modal state (STRiDE) separately. STRiDE1 and shows group-level statistical significance between the groups.

Next, ICA was applied to all 20 generated structural connectivity matrices to extract two independent components, representing simulated structural bases. The number of components (two) was selected using the elbow criterion.

To simulate dynamic functional network connectivity (dFNC), adjusted structural bases were first created by adding Gaussian variable (mean = 0, standard deviation = 0.1) to the two original structural bases. These adjusted bases were then used to simulate two groups of dFNCs. Group differences were introduced by further manipulating the adjusted structural bases. For each group, dFNCs were generated for 15 subjects by combining the adjusted bases over 50 time points with a random mixing matrix (mean = 0, standard deviation = 1). Subject-level variability was modeled by adding Gaussian noise (mean = 3, standard deviation = 3) to the estimated dFNCs.

For single-modal state estimation, we replaced the structural bases with functional ones in the proposed model. Functional bases were extracted by applying ICA to the generated dFNCs. These two functional bases, along with the dFNCs, were input to the connectivity-constrained ICA model to compute single-modal states for each subject. The average single-modal states are shown in Figure 2.c. In parallel, the original structural bases, along with the dFNCs, were used to compute multimodal states (i.e., simulated STRiDEs), shown in Figure 2.e.

Model performance was evaluated using a two-sample t-test on the estimated single-modal and multimodal states. Since group differences were embedded in both structural and functional data, we hypothesized that structural connectivity would enhance the detection of group effects. Additionally, we assessed the model’s robustness to noise—given the inherent noise in MRI data— to evaluate its stability under noisy conditions.

### 3. Real data

#### Participants/Imaging

Imaging data used in this study were obtained from the phase III of Functional Biomedical Informatics Research Network (FBIRN) dataset [35]. We used resting state functional MRI (rs-fMRI) and diffusion MRI (dMRI) images scanned from 311 age-gender-matched individuals, consisting of 162 healthy controls (HC) (45 females and 117 males), Average ± standard deviation (SD) age: 37.0 ± 11.0 and 149 participants with schizophrenias (SZ) (36 females 113 males), Avgerage ± SD age: 37.9 ± 11.5. Imaging data were collected on six 3T Siemens Tim Trio System and one 3T GE Discovery MR750 scanner at multiple sites, imaging details are in [36].

#### Preprocessing

The rs-fMRI images were preprocessed using the statistical parametric mapping (SPM12; https://www.fil.ion.ucl.ac.uk/spm/) toolbox in MATLAB 2020b according to the pipeline in [37]. This includes removing the first five fMRI time points, motion correction, slice timing correction, image registering to the standard Montreal Neurological Institute (MNI) space, spatial resampling (to 3 × 3 × 3 mm^3^ isotropic voxels), and spatial smoothing using Gaussian kernel with a full width at half maximum (FWHM) = 6 mm.

To preprocess the dMRI images, a similar procedure as in [38] was applied using the following steps. First eddy current and motion effects were corrected by motion-induced signal dropout detection based on b0 volumes and replacement method [39] through the eddy package (FSL 6.0; [40]). Then, we performed data quality check using an in-house-developed algorithm and visual inspection to exclude images with extreme head motion, signal drop out, or noise level [38]. After preprocessing, 274 of subjects remained for the future analysis.

#### Brain Network Connectivity

Next, we build two types of brain networks connectivity using the preprocessed rs-fMRI and dMRI images. To construct brain connectivity network, first intrinsic functional networks were determined using NeuroMark_fMRI_1.0 atlas [29], consisting of *N* = 53 replicable networks. To avoid overlapping, the ICNs were thresholded (values more than 3). The 53 networks are grouped into seven functional domains: subcortical (SC), somato-motor (SM), auditory (AUD), visual (VS), cognitive control (CC), default-mode (DM) and cerebellum (CB). Then, dynamic functional network connectivity (dFNC) was computed using the specific time courses of each network for individuals. To assess the dFNC a sliding window correlation approach was employed with a window size of 30 TRs (60 seconds) and 1 TR step size [1, 34]. Window was designed as a rectangular frame of 30 time points, convolved with a Gaussian kernel (σ = 3 TRs) to tap the window along the edges. By sliding the window across the time, pair-wise Pearson’s correlation was calculated in each time interval. The analysis yields a distinct functional network connectivity (FNC) at different time window for each subject.

To assess individual-level structural connectivity, voxel-wise diffusion tensor was first estimated using dtifit packages under FSL toolbox on preprocessed dMRI data. Whole-brain deterministic tractography was then, performed using the Camino toolbox [41] with 0.5 mm step size, 0.2 anisotropy threshold and 60° angular threshold. The resulting fiber tracts and fractional anisotropy (FA) maps were in the native space. FA maps were subsequently transformed to MNI space to obtain the NeuroMark_fMRI_1.0 atlas in native space via inverted spatial normalization using advanced normalization tools (ANTs) [42]. Finally, *N*×*N* structural network connectivity (SNC) matrices were generated for each subject by combining the normalized atlas with tractography results, where number of streamlines connecting each network pair defined the connection weights.

#### Statistical analysis on the STRiDE and TC

In this study, we performed a couple of statistical analysis to evaluate the performance of our model, identifying significant differences between healthy control and participants with schizophrenia, cognitive scores including speed of processing, attention vigilance, working memory, verbal learning, visual learning, reasoning problem solving and CMINDS composite), and symptoms scales such as PANNS positive and negative. A generalized linear model (GLM) was applied to the identified STRiDE, corresponding TC, their state vectors parameters and power. To control the effect of other influenced factors on the GLM analysis, we included four other independent variables in addition to the diagnosis of schizophrenia, cognitive scores and symptoms scales. The additional independent variables were age, gender, imaging site and head motion. To capture the significant cells, we considered the cells with FDR-corrected P-values bellow 0.05.

## Results

In this study, we applied a connectivity-constrained, data-driven ICA model, referred as STRiDE, to induce structural connectivity information on dynamic functional interactions in the brain. The performance of the proposed model was first evaluated using a simulation pipeline. We further assessed the model’s applicability by examining its performance on a real dataset and analyzing the effects of schizophrenia on multimodal brain states in comparison with healthy control subjects.

### Model Evaluation with Simulated Data: Validating the Integration of Structural and Dynamic Functional Connectivity

Figure 2 presents a hypothetical simulation illustrating the advantages of the proposed multi-modal model (STRiDE) compared to a single-modal approach. Panel (a) displays two representative connectivity patterns used as ground truth sources to generate the simulated data. The target area, highlighted in panel (a), indicates the manipulated connections designed to simulate group-level differences. Panel (b) shows the estimated structural bases derived from the simulation pipeline based on the two ground truth sources. Panel (c) presents the average of the single-modal states estimated using only functional data, and panel (d) shows the results of two-sample t-tests performed separately on each single-modal state. State 1 (imposed to show group differences) of the single-modal exhibits no significant statistical differences between groups (FDR-corrected P-value>0.05). In contrast, panel (e) shows the average of the multi-modal states estimated by STRiDE, which integrates both structural and functional data. The corresponding statistical tests in panel (f) reveal significant group-level differences (FDR-corrected P-value<0.05), in simulated STRiDE 1. These findings demonstrate the enhanced sensitivity and statistical power of STRiDE in detecting group differences through the integration of multimodal information.

A comparison the resiliency of the single-modality and multi-modality model for the noise is presented in Figure 3, for different noise levels. In Figure 3, we show the number of significant connections identified by single-modality and multi-modality models under varying noise levels. Results illustrate that the multi-modal model (STRiDE) consistently detects a greater number of significant connections compared to the single-modal approach based on the ground truths. While the single-modal model shows more spread-out results with greater variability and fewer significant connections overall, compared to the multi-modal model (simulated STRiDE) with less variability and higher significant connections across noisy conditions. These findings suggest that STRiDE is more robust and sensitive in detecting true group-level effects, even under noisy conditions, highlighting its advantage in leveraging both structural and functional information to enhance the model stability. We note that this simulation is meant to demonstrate the approach, rather than provide a comprehensive evaluation of all possible scenarios.

**Figure 3.**
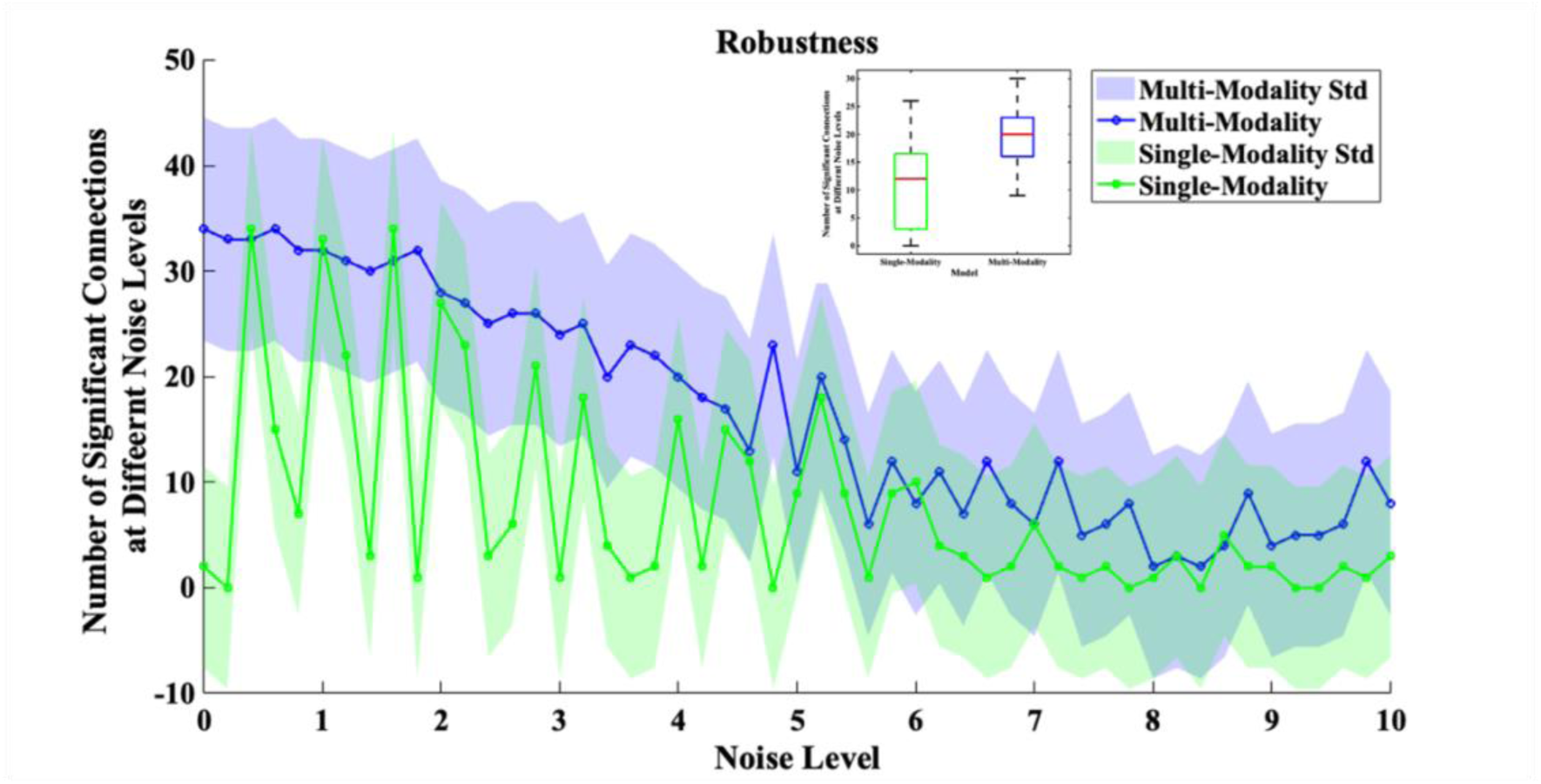
Comparison of the number of significant connections identified by single-modality and multi-modality (STRiDE) models under varying noise levels at the target area. The multi-modal model consistently detects more significant connections with reduced variability, demonstrating greater sensitivity and robustness to noise compared with the single-modal approach.

### Real data results

#### Model fitness to structural and functional information

To determine the amount of structural information captured by estimated multimodal states (STRiDE), we estimate similarity (Pearson’s correlation) of the STRiDEs with structural bases. Our estimation illustrates strong correspondence between the estimated STRiDE and the reference structural connectivity bases (Bases), shown in Figure 4. A higher value along the diagonal (Figure 4.a), indicate a substantial degree of similarity between each estimated STRiDE and its identically related structural basis, ranging from 0.48 to 0.61 (Figure 4.a). Specifically, all STRiDEs demonstrate a significant high correlation (P-value<0.05; ρ = [0.48 – 0.61]) with corresponding structural bases. This consistent pattern of high diagonal values underscores the model’s effectiveness in leveraging structural prior information to accurately reconstruct the underlying connectivity patterns, which has observed clearly in P-values as well, as -*log_10_(P-value)*sign(T-value)* is shown Figure 4.b. The relatively low off-diagonal elements further support this strong accordance, indicating minimal similarity between an estimated STRiDE and any non-corresponding structural basis. These results highlight the robust ability of our multimodal approach to successfully disentangle the structural bases, yielding estimated connectivity patterns that closely mirror the provided structural prior information across all identified STRiDEs.

**Figure 4.**
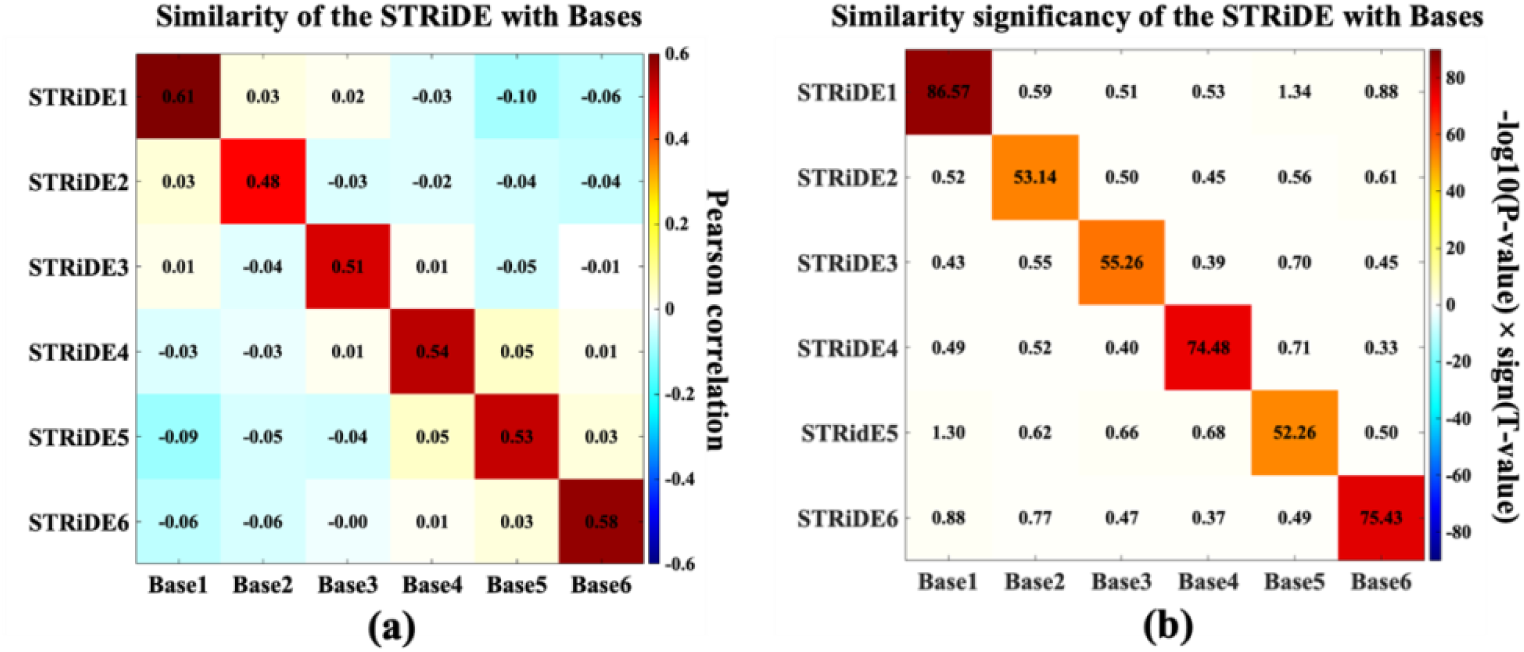
Illustrates the estimated STRiDEs similarity (Pearson’s correlation) and their similarity significancy with structural information. In (a) average similarity (Pearson’s correlation) of the six estimated multimodal states (STRiDEs) with structural Bases, while p-values of the estimated correlations shown in (b) as (-log_10_(P-value)×sign(T-value). Overall, significantly correlated (P-value<0.05) STRiDEs with their corresponding Bases represented the independently of each STRiDEs and highlights the existing of distinguished reoccurring states in brain, supported by structural and functional information.

Statistical fitness (R^2^) for the linear regression of estimated STRiDEs on dynamic functional information determines the amount of functional information explained by structural. It was examined using generalized linear model (GLM) while controlling the effect of age, gender, site and motion (mean framework displacement).

It showed that on average less than one third (**mean** ± **std = 0.27** ± **0.05**; **maximum-minimum = 0.15 – 0.62**) of the dynamic functional connectivity information was included in the STRiDEs. In addition, state-specific R^2^ estimation for each STRiDEs, followed by a two-sample t-test analysis reflected that the estimated R^2^ are significantly (P-value<0.05) associated with SZ, specifically, it showed lower values for the SZ in STRiDE1, STRiDE4 and STRiDE5, while STRiDE6 was significant higher in SZ subjects.

### Structural bases: relating brain structure and dynamic function to characterize reoccurring brain states

To identify the reoccurring brain states, we constrained the brain dynamic functional information during rest via a multi-modal optimization which leverages structural bases. These structural bases, shown in Figure 5, are obtained from independent components derived from the individual-level structural connectivity and used as a reference to inform the whole brain dynamic functional information at connectivity domain. The ICA of structural connectivity revealed six distinct bases (based on elbow criteria) that encapsulate spatially organized patterns of variability across the brain’s structural architecture (Figure 5). These bases reflect modular connectivity features across canonical functional networks, including unimodal networks/regions [43] (e.g., sensorimotor (SM), visual (VS)) and trans-modal regions [43] (e.g., default mode (DM), cognitive control (CC)). Structural Base1 exhibits dominant intra-network connectivity within unimodal networks and mild interactions with higher-order networks, suggesting a foundational structural motif. In contrast, Bases3 and Base5 show increased connectivity involving trans-modal and cerebellar networks, indicative of more integrative, higher-level structural organization. Base6 reveals focused cross-network connections, particularly linking networks of the VS and CC domains, consistent with intermediate-level processing. This gradation across bases—from localized, low-level sensorimotor patterns to distributed, high-level trans-modal integration—suggests that the ICA decomposition captures a hierarchical structure of brain organization embedded in the structural connectome.

**Figure 5.**
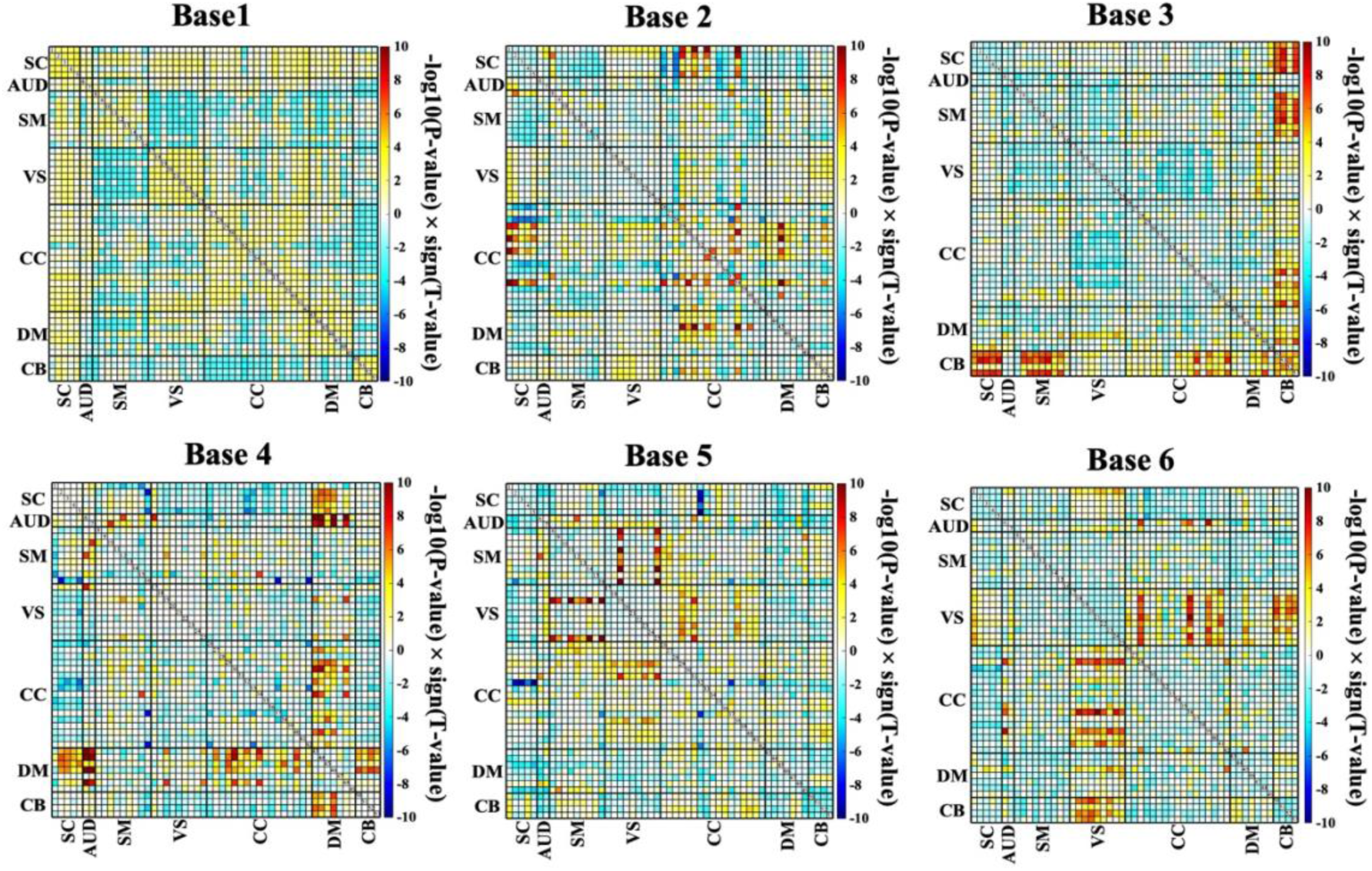
The connectivity pattern of the GICA-derived structural connectivity bases (1–6). Each matrix depicts a set of independent structural connectivity patterns observed across a group of subjects, ordered by intrinsic functional connectivity networks: SC, AUD, SM, VS, CC, DM, CB. The components capture distinct intra- and inter-network patterns, reflecting modular and hierarchical organization, from unimodal segregation to trans-modal integration.

### Structural connectivity-informed ICA model exhibits unique hierarchical multimodal brain states (STRiDEs)

The proposed multimodal model allowed us to estimate six structurally informed and temporally varying multimodal connectivity patterns-hereafter referred to as STRiDEs that capture dynamic functional interactions shaped by underlying structural motifs (Figure 6). Each multimodal states reflects a distinct connectivity profile aligned with specific structural bases. Notable, some states exhibit strong intra-network coherence, such as SM and VS in STRiDE1, while others highlight cross-network interactions such as VS-CC (in STRiDE6), DM-CB (in STRiDE 3), consistent with higher-order integrative processes. The use of structural priors yielded more anatomically plausible and interpretable states compared to unconstrained models, preserving both modular integrity and inter-network dynamics [8].

**Figure 6.**
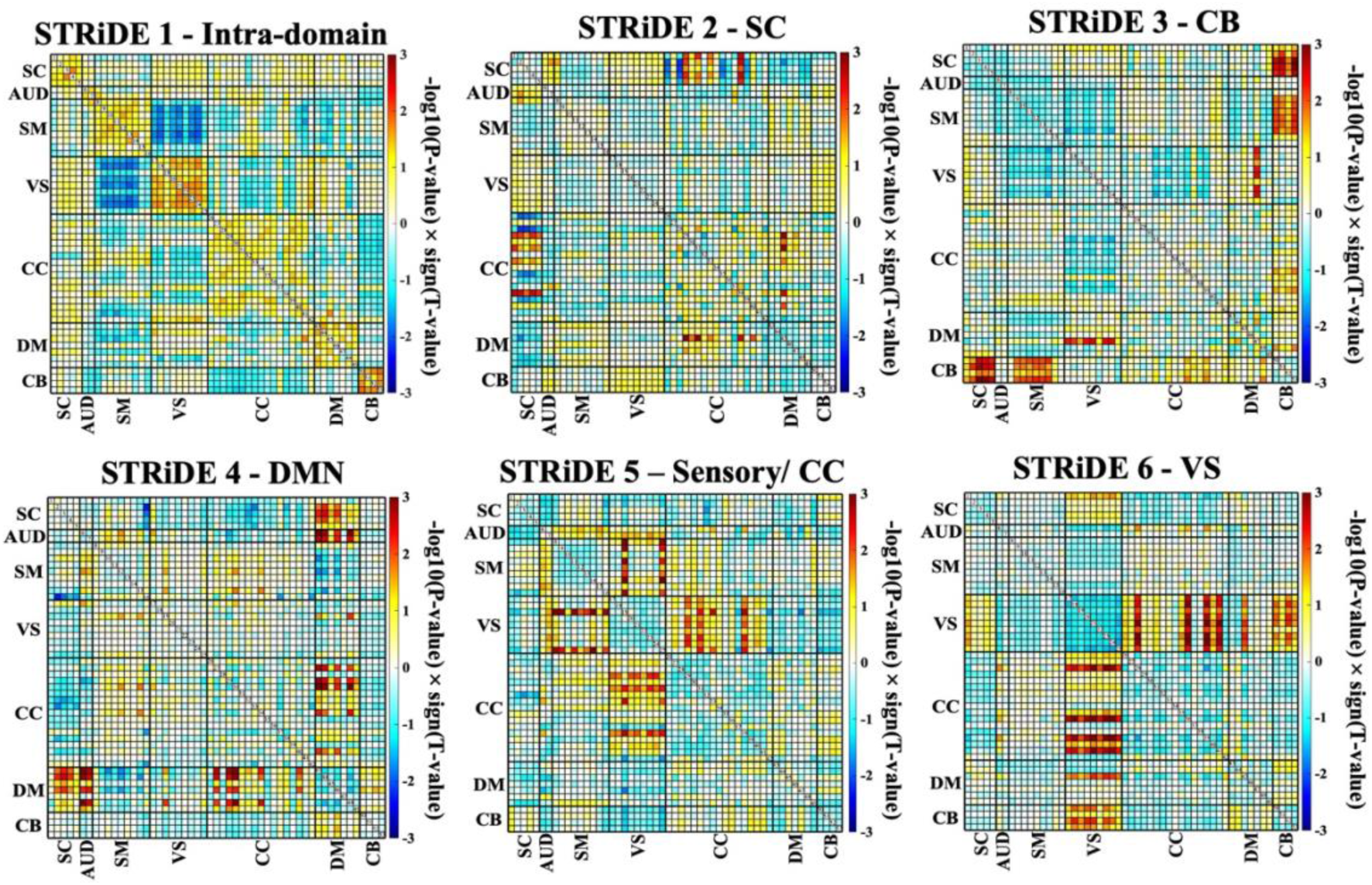
displays the connectivity pattern of the structurally constrained multimodal connectivity states (STRiDE). Six distinct multimodal states, STRiDEs, were estimated using structural connectivity bases as priors within a regularized multimodal data-driven model. Each matrix represents a different dynamic functional connectivity state shaped by the underlying structural architecture, across different functional domains (SC, AUD, SM, VS, CC, DM, CB). The STRiDEs reveal unique intra- and inter-network connectivity patterns, capturing structural-informed dynamics that reflect modular, integrative, and hierarchical brain organization.

The observed multimodal states reflect a spectrum of brain network configurations ranging from localized, unimodal segregation to distributed, trans-modal integration. This alignment with the structural gradient supports the role of anatomical architecture in shaping functional dynamics. Together, these results demonstrate that structurally informed multimodal modeling can disentangle meaningful and hierarchically organized patterns of time-varying connectivity.

### Multimodal States (STRiDE) Reveal State-specific Dysconnectivity Patterns Correlated with brain features in participants with SZ

Using a statistical analysis on STRiDEs across group of HC and participants with SZ, shows a significant difference between connectivity pattern of the HC and SZ. For the statistical analysis we applied a general linear model (GLM), adjusting for potential confounding variables including age, gender, head motion, and imaging site. State-specific connectogram plots in Figure 7 visualized the significant connections (FDR-corrected P-value <0.05) per estimated STRiDE, each nodes are different brain regions divided into seven domains (color-specific); subcortical (SC), auditory (AUD), sensory motor (SM), visual (VS), cognitive control (CC), default mode (DM), and cerebellum (CB). The results reveal distributed pattern of dysconnectivity across intra- and inter-domain interactions. Notably, certain states (e.g., STRiDE 1 and STRiDE 4) exhibited widespread alterations predominantly in sensory (SM, AUD, VIS) and cognitive-control (CC) domains, whereas others (e.g., STRiDE 3 and STRiDE 6) highlighted focal distributions within specific domains such as the DMN and subcortical. In contrast, traditional single modality (functional-only) approaches often report dysconnectivity within and between high-order domains and frontoparietal control domain, often manifesting as reduced within-domain cohesion and disrupted cross-domain integration [34, 44]. These observations suggest that dynamic connectivity alterations in SZ are state-dependent, with distinct STRiDEs capturing different facets of disease-relevant functional disruption, aligned with known structural vulnerabilities in schizophrenia-such as connectivity changes in and between sensory and trans-modal domains.

**Figure 7.**
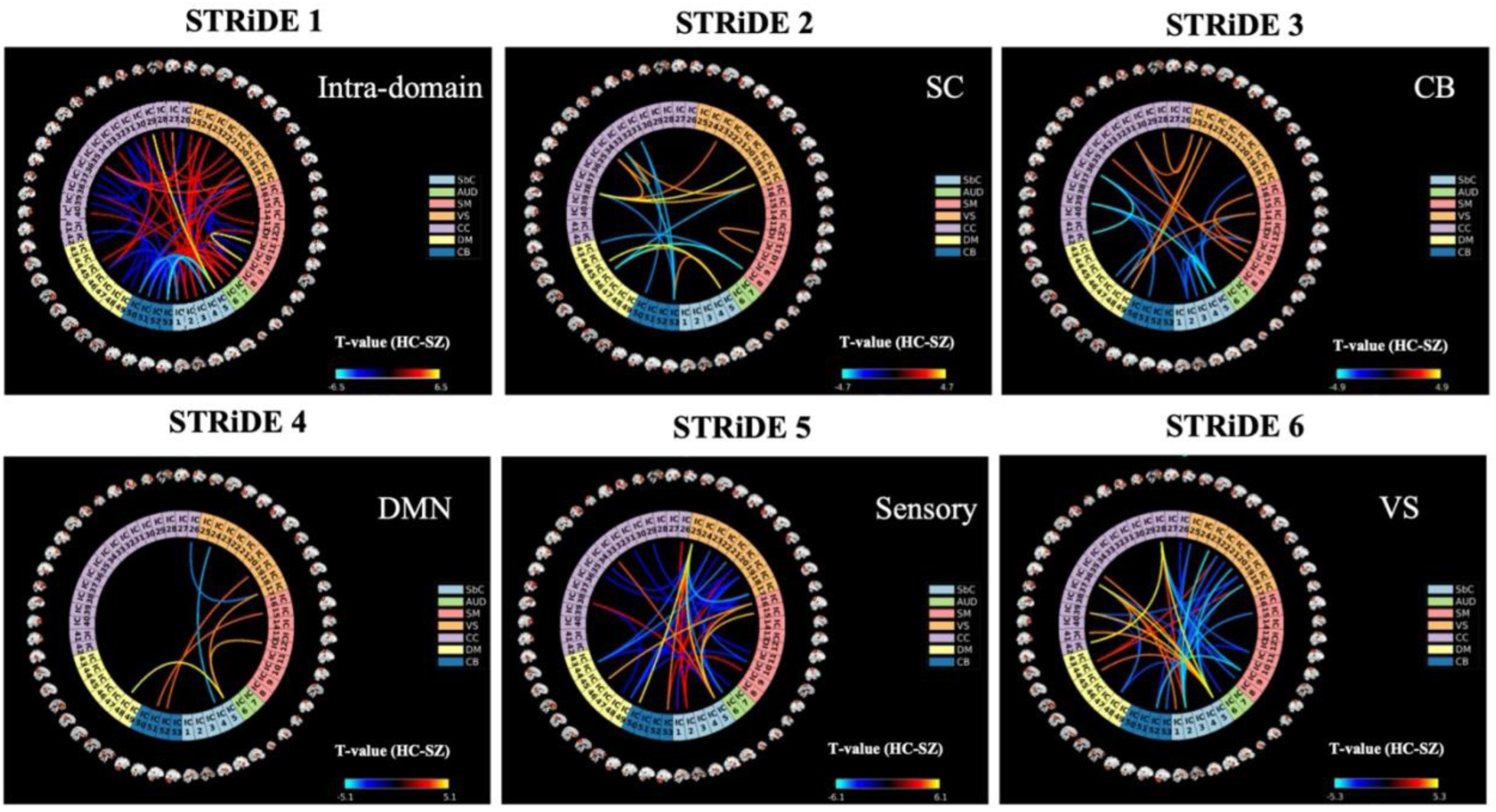
illustrated statistical analysis, using a generalized linear model (GLM) on the STRiDEs. It shows the significant connection (FDR-corrected P-value < 0.05) observed in multimodal states (STRiDEs) which are different between healthy control (HC) and schizophrenia (SZ). Each state-specific connectogram reveals the most significant dysfunctionality observed in Intra-domain, Sensory-CC related and VS related domains between the HC and SZ.

Incorporating structural priors into functional estimation results in spatially constrained dysconnectivity maps that are more biologically interpretable. This is particularly beneficial in structurally disrupted disorders such as schizophrenia (SZ), where the findings are more reliably linked to underlying neuropathology [45]. In addition, providing multiple modality information for brain state estimation can strengthen the findings of the underlying mechanism of brain as in our results, offer improved sensitivity to subtle group differences which is often overlooked in functional-only models. Importantly, the significant group differences observed here thus reflect how structure-constrained dynamics can disentangle complementary aspects of functional dysconnectivity in schizophrenia, aligning with the idea that distinct structural motifs underlie state-specific vulnerability. This supports the utility of the STRiDE model in bridging the structural and functional dimensions of neuropsychiatric disorders.

Furthermore, statistical analysis on estimated STRiDEs, was applied to investigate association of the connectivity disruption with seven different cognitive scores, including speed of processing, attention vigilance, working memory, verbal learning, visual learning, reasoning problem solving and CMINDS composite as presented in Figure 8. The analysis reveals that there are significant (FDR-corrected P-value<0.05) differences in altered interactions related to four of the cognitive scores (attention vigilance, working memory, visual learning and CMINDS composite) between HC and SZ subjects. Scores differences were state-specific, where each states shows differences for one score, except the STRiDE 6.

**Figure 8.**
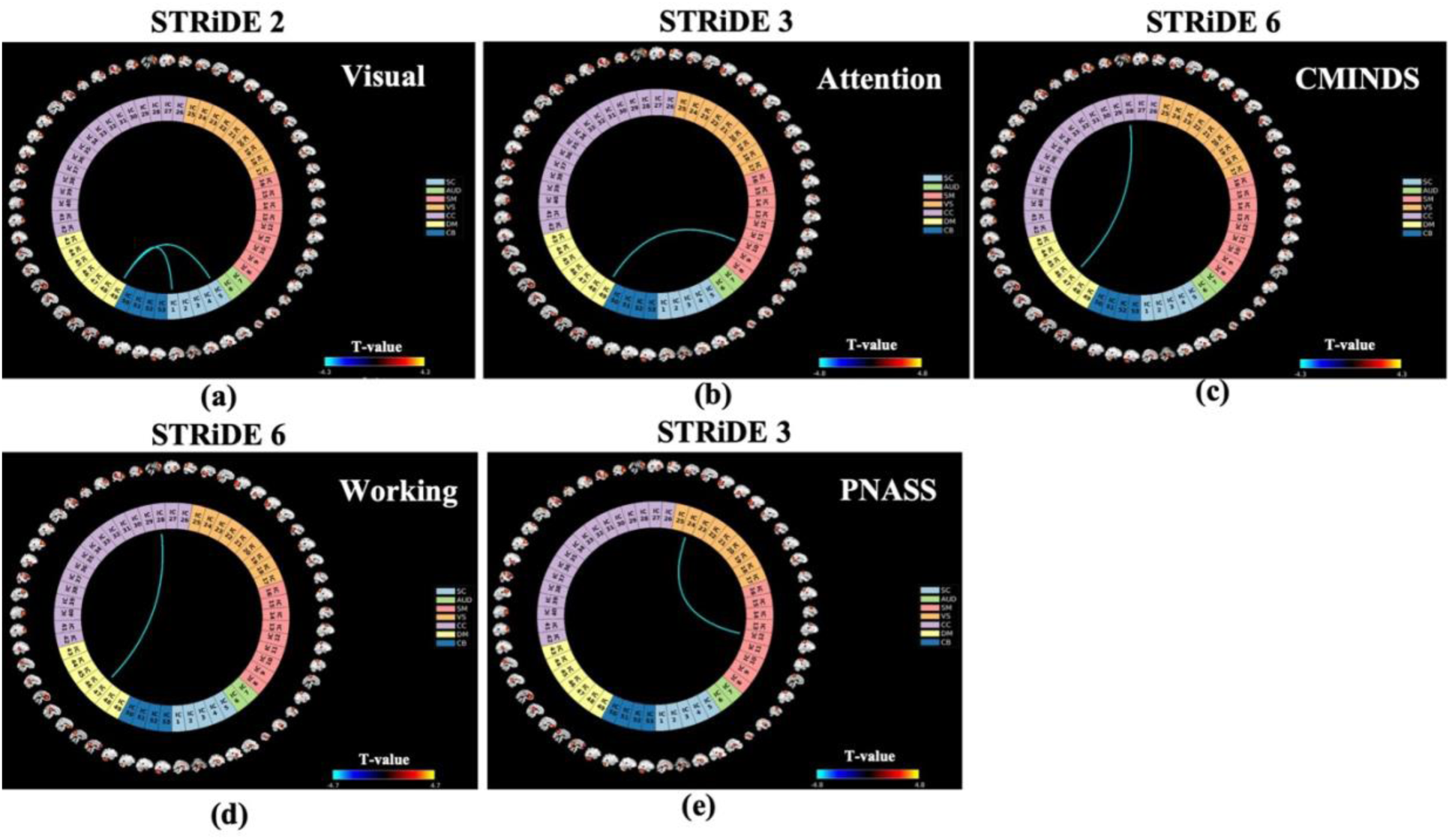
State-specific statistical analysis (GLM) of STRiDE connectivity in relation to cognitive performance and clinical symptoms. Significant associations were found in three STRiDE states with cognitive measures including visual learning, attention/vigilance, working memory, and the CMINDS composite score, as well as with the PANSS positive symptom scale. The GLM was adjusted for age, gender, head motion, group, and imaging site. Panels (a–d) show significant effects (FDR-corrected p < 0.05) of the four cognitive scores on inter-network connectivity, particularly between the DM-SC, DM-SM, and DM-CC domains. Panel (e) highlights a significant symptom-related connectivity difference within the schizophrenia group between the VS and SM networks. These findings underscore the role of structural information in capturing connectivity changes, especially in trans-modal domains (DM, CC) as well as unimodal sensory domains in schizophrenia.

Figure 8.a-b, shows that in STRiDE 2, there are two significantly different connections but negative; between DM and SC domain for the visual learning score, while only one connection between DM and SM was significant in STRiDE 3 with negative color for the attention vigilance scores. STRiDE 6 shows significant differences regarding two different cognitive scores, in Figure 8.c-d. Interestingly, connection alteration between the CC and DM shows significant different (FDR-corrected P-value<0.05) between the HC and SZ related to the working memory and CMINDS composite. These findings suggest that the functional organization of the brain, as captured by these multimodal states, plays a crucial role in attention vigilance, working memory, visual learning and CMINDS composite abilities, with specific inter-regional communication patterns within STRiDEs 2, STRiDE 3 and STRiDE 6, showing a strong association with individual differences in those three cognitive scores. Furthermore, symptom-related dysconnectivity in estimated STRiDEs, illustrated association with positive symptoms in STRiDE 3 for SZ.

Figure 8.e displayed the statistical analysis results that reveals the existence of only one significant connection (FDR-corrected P-value<0.05) differs within the SZ cohort, as observed between visual (VS) and sensory motor (SM) domain. It is showing significant association with PANSS in the SZ group. The connectogram highlights connections where increased or decreased connectivity is significantly related to the severity of positive symptoms (e.g., hallucinations, delusions). These findings suggest that STRiDE 3 may reflect a symptom-relevant state, capturing transient network dynamics linked to the expression of positive clinical features, and underscores the value of multimodal models in revealing state-specific symptom connectivity profiles.

### Temporal Interplay Among Multimodal Brain States STRiDEs

Building on the spatial architecture and occurrence dynamics of the STRiDEs, we next examined the temporal interrelationships among their time courses (TC). Figure 9.a illustrates the average similarity (Pearson’s correlation) matrix of all STRiDE TCs, revealing both positive and negative associations across states—most notably, strong negative correlations involving TC5 with TC2 and TC3, and TC3 with TC1. The corresponding *–log_10_(p-values*), in Figure 9.b highlighting the statistical robustness of these correlations using two-sample t-test, showing the nonsignificant value in the off-diagonal cells (similarity of each TC with other remaining TCs). Panel (c) presents group differences (HC vs. SZ) in TC similarity *(-log_10_(FDR-corrected P-value)*×*sign(T-value))*, identifying altered relationships particularly between TC3 and other states in schizophrenia. These findings suggest dysregulated temporal coordination among brain states in SZ, potentially reflecting disrupted dynamic integration in functional brain networks.

**Figure 9.**
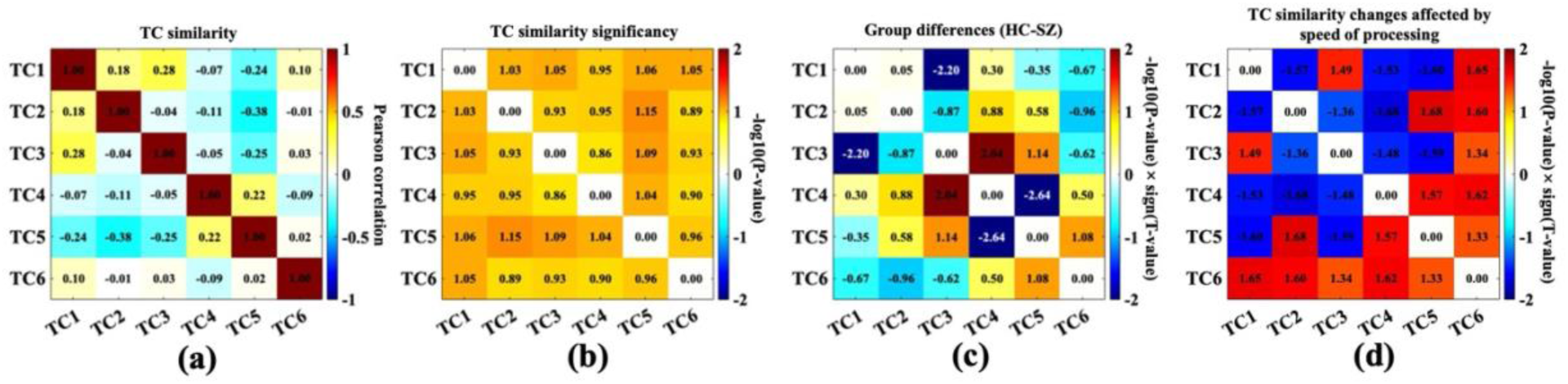
displays the temporal similarity (Pearson’s correlation) of the corresponding time courses (TCs) with each STRiDEs and the association of the correlations with SZ and the cognitive score related to the speed of processing using statistical analysis. In panel (a), the average similarity between the TCs reveals distinct patterns of correlation and anti-correlation where, shows significant (two-sample t-test; P-value<0.05) similarity between the identical TCs. Panel (b) is the *-log_10_(P-value)* of the estimated P-values. In addition, TC similarity association with SZ, was explored by a GLM analysis, (*-log_10_(FDR-corrected P-value)*×*sign(T-value)*) displayed in panel (c), with significant alterations indicating disrupted temporal coordination among specific STRiDEs in SZ. In panel (d), GLM analysis highlights the TC similarity correlation with speed of processing score, (*-log_10_(FDR-corrected P-value)*×*sign(T-value)* shown in panel (d)). These findings collectively illustrate how speed of processing influences the dynamic interplay among multimodal brain states.

Moreover, the strong negative correlation between TC5 and TC2/TC3 aligns with the spatial antagonism observed between these STRiDEs in their multimodal connectivity profiles—where STRiDE5 emphasizes dysconnectivity in cognitive control regions, while STRiDE2 and STRiDE3 are more associated with sensory and default-mode domains. Results shows that the temporal relationships are not random but instead reflect underlying spatial dependencies among the STRiDEs—states with overlapping or complementary spatial topographies tend to exhibit stronger temporal similarity. This overlay of temporal correlation and spatial architecture highlights the hierarchical and structured nature of the brain’s dynamic organization.

Further investigation of the brain connectivity disruption in schizophrenia regarding the effect of the cognitive score and symptoms on the temporal interplay of the multimodal brain states, while considering the speeding of processing score between the HC and SZ, shown in Figure 9.d. Positive values (warmer colors) indicate that higher speed of processing score is associated with greater similarity in the temporal dynamics of the corresponding states, while negative values (cooler colors) suggest the opposite. Interestingly, the analysis reveals significant (*-log_10_(FDR-corrected P-value)*×*sign(T-value))* associations between speed of processing and time course similarity for several STRiDEs pairs. For instance, a strong positive association was observed between speed of processing and the similarity of TC1 and TC6, while a significant negative association was found for TC2 and TC4. However, the interplay of the STRiDE 6 is positively correlated with temporal similarity of all other STRiDEs. Other STRiDEs mostly interact negatively with others except the for the TC1 and TC3, TC2 and TC5 and TC4 and TC5. These results suggest that an individual’s speed of processing ability influences the degree to which different multimodal brain states (STRiDE) exhibit synchronized or desynchronized temporal fluctuations. Given that TC6 is related to the visual system, indicates the link between the visual system and speed of processing, is another confirmation on the previous study [46] showing that visual attention and processing speed are related, and that visual training can improve processing speed. Overall, it is highlighted that an individual’s speed of processing ability influences the degree to which different multimodal brain states exhibit synchronized or desynchronized temporal fluctuations.

Total power of the estimated TCs, captures significant (FDR-corrected P-value<0.05) differences between the HC and SZ subjects for the TC1, TC4, TC5 and TC6, comes in Table 1. In accordance with findings on the connectivity pattern of the STRiDEs, we observed the differences in TCs related to the Intra-domain activity, DM domain activity, sensory and CC domain activity and VS domain activity. TC1, TC4 and TC5, show positive significant power differences, which indicate the STRiDEs1, STRiDE4 and STRiDE5 are mostly observed in HC than the SZ. In contrast, TC6 with negative significant power supports the higher occurrence of STRiDE6 in SZ compared to the HC. Results, reveals sensory domains, particularly VS is an important brain functional region in SZ cohort. Interestingly, additional statistical analysis on the power regarding cognitive scores, shows that the visual learning and reasoning problem solving scores significantly were affected in the TC4, TC5 and TC6. TC6 was positively correlated in visual learning, indicating that existence of more STRiDE6 increase the visual learning scores, which has highly occurred in healthy controls. Similarly, the TC4 was positively significant for the reasoning problem solving. Results highlight the importance of the default-mode related and visual related STRiDEs in increasing those scores. Visual learning scores was negatively correlated with TC4 (related to the default mode related STRiDE). Moreover, CMIND composite score was negative significant power for the TC5 (sensory related STRiDE). Other cognitive scores and symptoms show no significant power value.

**Table 1.**
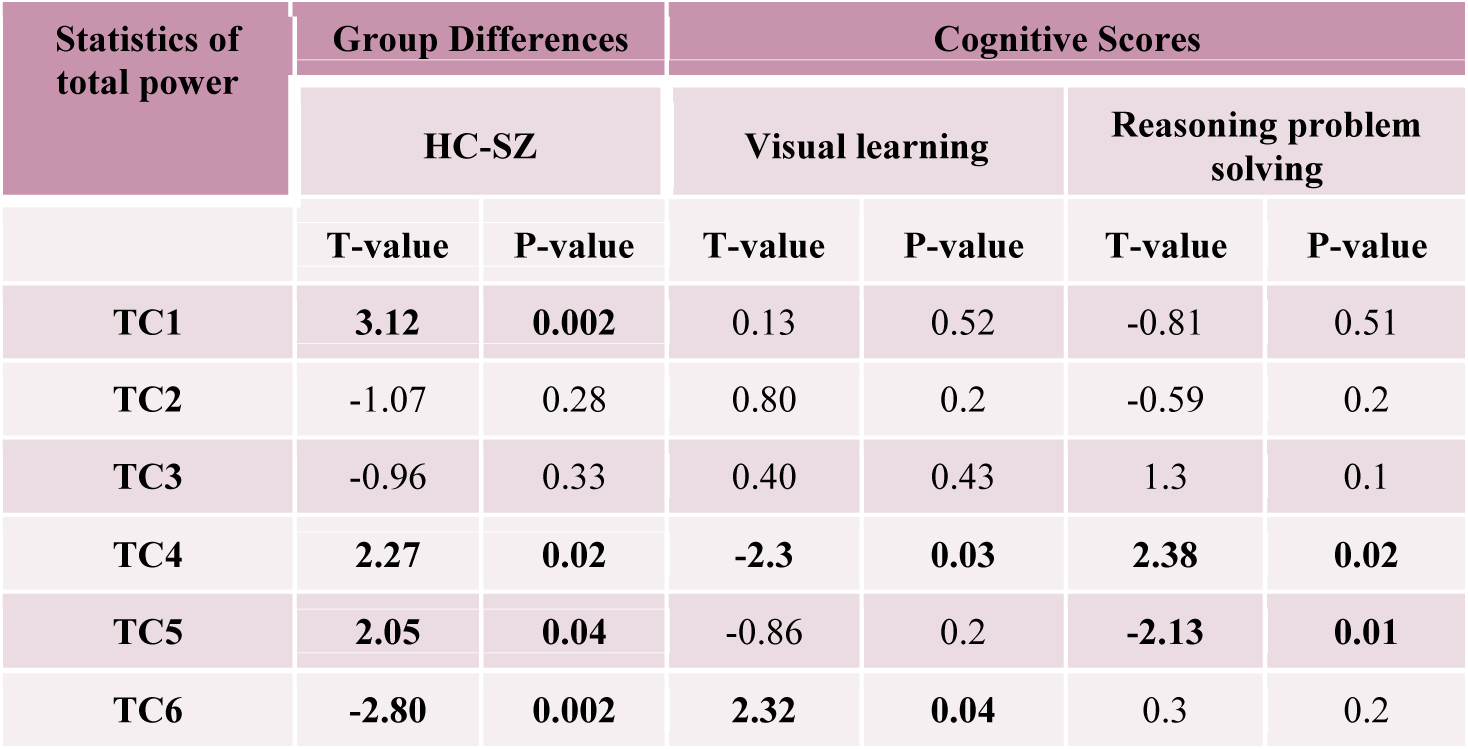
illustrates the statistical analysis (GLM analysis) on the estimated total power regarding two significant cognitive scores. Specific TCs were significantly different for the visual learning and reasoning problems.

| Statistics of total power | Group Differences |  | Cognitive Scores |  |  |  |
| --- | --- | --- | --- | --- | --- | --- |
|  | HC-SZ |  | Visual learning |  | Reasoning problem solving |  |
|  | T-value | P-value | T-value | P-value | T-value | P-value |
| TC1 | 3.12 | 0.002 | 0.13 | 0.52 | -0.81 | 0.51 |
| TC2 | -1.07 | 0.28 | 0.80 | 0.2 | -0.59 | 0.2 |
| TC3 | -0.96 | 0.33 | 0.40 | 0.43 | 1.3 | 0.1 |
| TC4 | 2.27 | 0.02 | -2.3 | 0.03 | 2.38 | 0.02 |
| TC5 | 2.05 | 0.04 | -0.86 | 0.2 | -2.13 | 0.01 |
| TC6 | -2.80 | 0.002 | 2.32 | 0.04 | 0.3 | 0.2 |

### Temporal Dynamics of Structurally Constrained Multimodal Brain States (Strides) Highlights the Hierarchical Characteristics of the STRiDE output

The state vector analysis provides a detailed characterization of the temporal dynamics underlying the six identified multimodal brain states (STRiDEs) in Figure 10. STRiDE1 emerged as the most dominant state, exhibiting the highest frequency of occurrence, occupancy rate, and mean dwell time, indicating a highly stable and recurrent network configuration. In contrast, the other STRiDEs appeared less frequently and were sustained for shorter durations. The transition probability matrix further supports this, with high diagonal values reflecting strong within-state persistence—particularly for STRiDE1, STRiDE2, STRiDE4, STRiDE5, and STRiDE6—and most frequent transitions observed from STRiDE1 to STRiDE3 and STRiDE6. The average number of transitions per subject was **3.6 ± 3.1** (**Mean ± Std**), suggesting moderate temporal switching. Results are aligned with previous single modal modes, however, the temporal dominance of STRiDE1 and transient nature of other states reflect the brain’s hierarchical organization, with stable activity in lower-order unimodal networks/regions and flexible transitions in higher-order trans-modal regions. This pattern, supported by structurally constrained estimation, aligns with earlier structural base findings and known cortical hierarchies [11].

**Figure 10.**
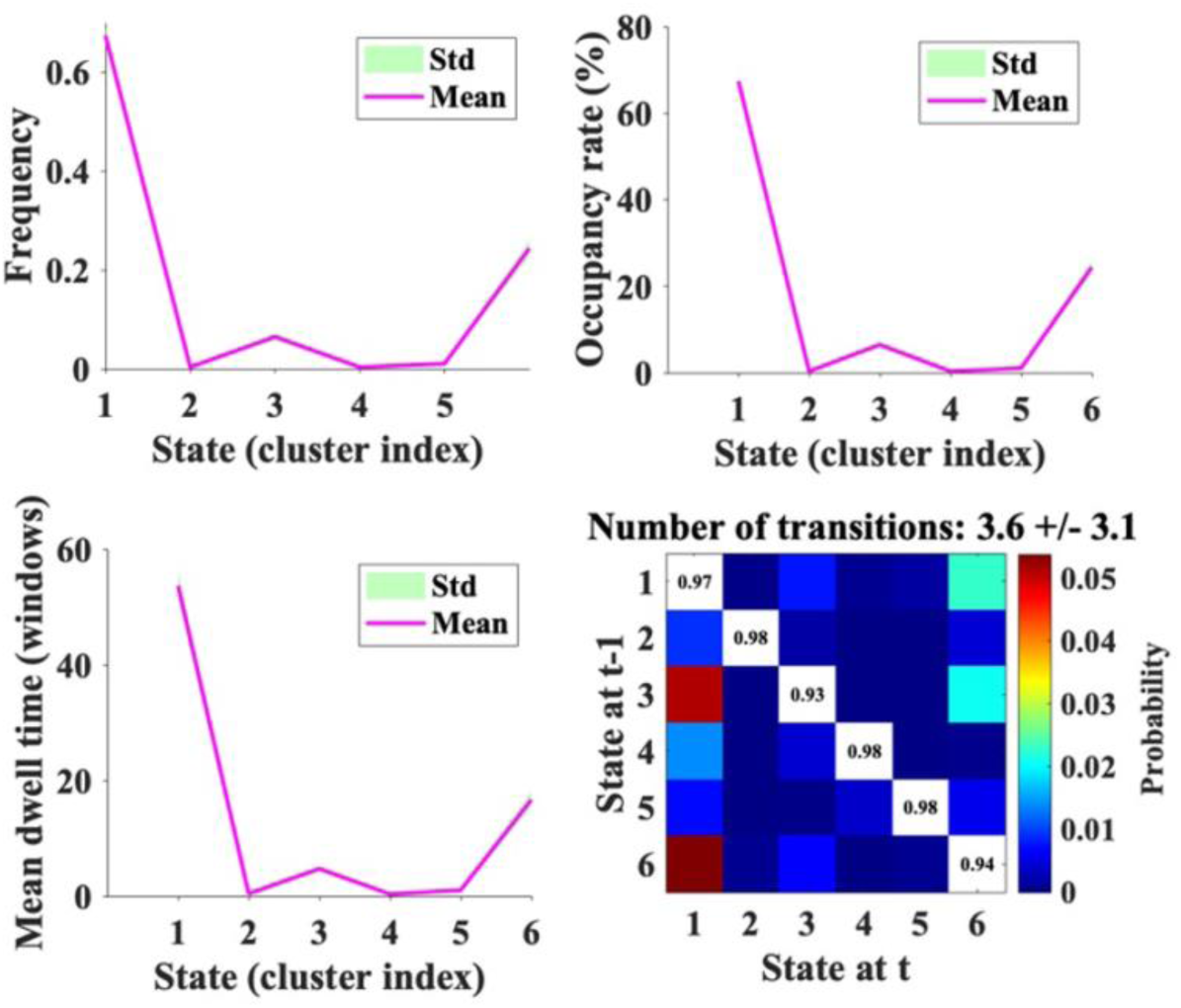
Temporal dynamics of multimodal brain states (STRiDEs) derived from structurally constrained estimation model. The top-left panel illustrates the frequency of occurrence for each of the six STRiDEs. While, occupancy rate is displayed on the top-right panel, indicating the percentage of time the system spends in each STRiDE. The average duration the system remains in each STRiDE before transitioning, is the mean dwell time which the bottom-left panel shows it. State vector analysis reveals that STRiDE1 is the most frequent, stable, and long-lasting state, while the other states appear less frequently and with shorter dwell times. The bottom-right panel presents the STRiDE transition matrix, depicting the probability of transitioning between STRiDEs. The transition matrix indicates high within-state stability and selective transitions, particularly from STRiDE1 to STRiDEs3 and STRiDE6. These patterns reflect underlying cortical hierarchies, with structurally informed estimation enhancing the alignment of dynamic states with unimodal-to-trans-modal organization.

In addition, statistical analysis on the state parameters, using the GLM model, shows significant differences (Table 2) for the mean dwell time and occupancy rate for STRiDE1, STRiDE2, STRiDE3 and STRiDE6 between the HC and SZ. Working memory scores were highly tied to the Mean dwell time over STRiDE1, showing positively affecting the working memory. For the verbal learning, it is positively related to the STRiDE3, while negatively to the STRiDE5. On the other hand, occupancy rate was significantly affected CMINDs composite score over STRiDEs 1, 5 and 6 (positively with the STRiDE6, and negatively with STRiDE1 and STRiDE5). In addition, STRiDE1, STRiDE2, STRiDE3 and STRiDE6 were significantly affected by the SZ. particularly, STRiDE1 was decreased in SZ.

**Table 2.**
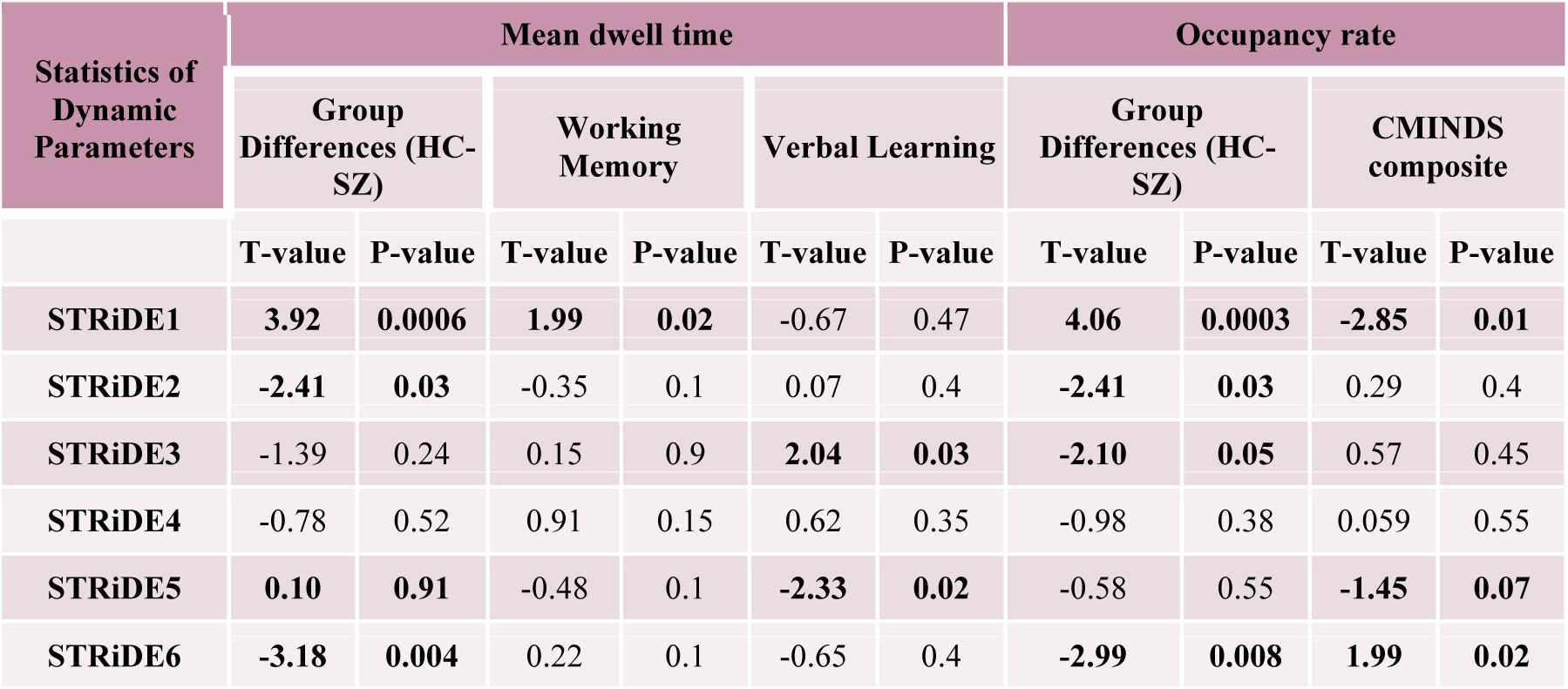
illustrates the statistical analysis (GLM model) on the estimated dynamic parameters regarding group differences and significant cognitive scores. Specific STRiDEs were mostly significant for the Mean dwell time and occupancy rate.

| Statistics of Dynamic Parameters | Mean dwell time |  |  |  |  |  | Occupancy rate |  |  |  |
| --- | --- | --- | --- | --- | --- | --- | --- | --- | --- | --- |
|  | Group Differences (HC-SZ) |  | Working Memory |  | Verbal Learning |  | Group Differences (HC-SZ) |  | CMINDS composite |  |
|  | T-value | P-value | T-value | P-value | T-value | P-value | T-value | P-value | T-value | P-value |
| STRiDE1 | 3.92 | 0.0006 | 1.99 | 0.02 | -0.67 | 0.47 | 4.06 | 0.0003 | -2.85 | 0.01 |
| STRiDE2 | -2.41 | 0.03 | -0.35 | 0.1 | 0.07 | 0.4 | -2.41 | 0.03 | 0.29 | 0.4 |
| STRiDE3 | -1.39 | 0.24 | 0.15 | 0.9 | 2.04 | 0.03 | -2.10 | 0.05 | 0.57 | 0.45 |
| STRiDE4 | -0.78 | 0.52 | 0.91 | 0.15 | 0.62 | 0.35 | -0.98 | 0.38 | 0.059 | 0.55 |
| STRiDE5 | 0.10 | 0.91 | -0.48 | 0.1 | -2.33 | 0.02 | -0.58 | 0.55 | -1.45 | 0.07 |
| STRiDE6 | -3.18 | 0.004 | 0.22 | 0.1 | -0.65 | 0.4 | -2.99 | 0.008 | 1.99 | 0.02 |

In Figure 11, statistical analysis on the transition matrix was displayed, transition from STRiDE1 (Intra-domain related) to STRiDE1, STRiDE2 (SC related) to STRiDE1 (Intra-domain related), STRiDE2 (SC related) to STRiDE2 (SC related), STRiDE4 (DM related) to STRiDE1 (Intra-domain related) and STRiDE5 (sensory/CC related) to STRiDE5 (sensory/CC related) was significantly different between the HC and SZ. Verbal learning association with transition matrix (Figure 11.b) reveals highly significant transition from STRiDE6 (VS related) to STRiDE4 (DM related), STRiDE1 (Intra-domain related) to STRiDE1 (Intra-domain related) and STRiDE1 (Intra-domain related) to STRiDE6. Transition from STRiDE2 (SC related) to STRiDE4 (DM related) and STRiDE5 (sensory/CC related), STRiDE3 (CB related) to STRiDE4 (DM related), STRiDE4 (DM related) to STRiDE2 (SC related), STRiDE4 (DM related) to STRiDE5 (sensory/CC related) and STRiDE5 (sensory/CC related) to STRiDE2 (SC related) are nonsignificant. In contrast, reasoning problem solving is highly correlated with only one transition from STRiDE5 to STRiDE5 (sensory/CC related), shown in Figure 11.c. All the panels in Figure 11 show *-log_10_(P-value)* × *sign(T-value)* at each matrix.

**Figure 11.**
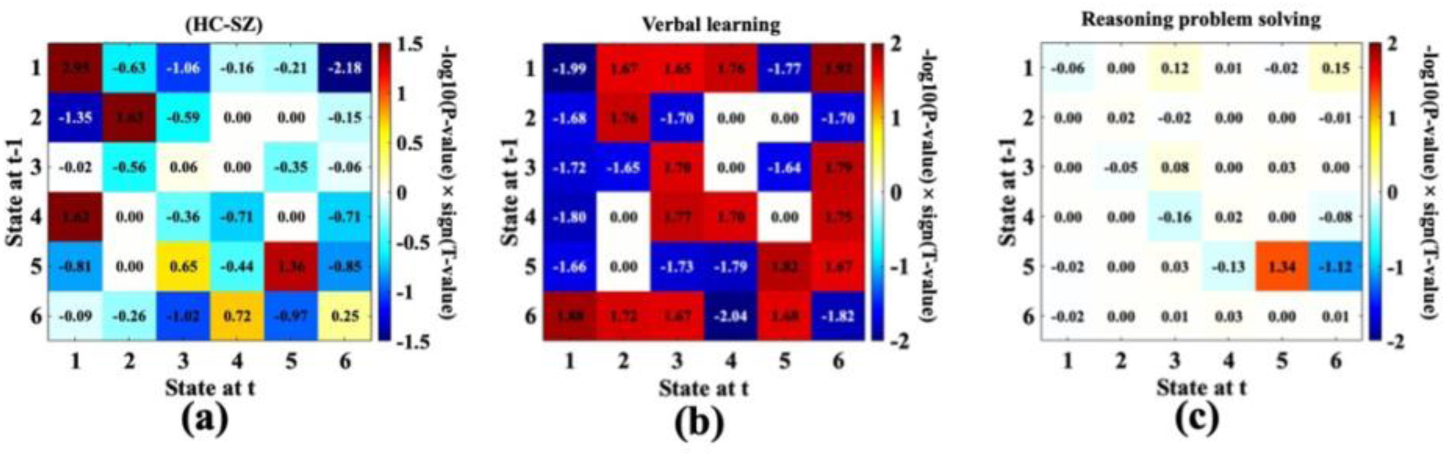
shows statistical analysis using GLM analysis on transition matrix investigating the group differences between HC and SZ in (a), verbal learning in (b), and reasoning problem solving in (c). Values are -log_10_(P-value)×sign(T-value). It highlights that the transition of STRiDEs related to interactions within the Intra-domain and trans-modal domains were significantly correlated with SZ, while verbal learning was significantly affected by SZ over most of the STRiDE transitions, specifically increasing of the transitions from intra-domain related STRiDE to other STRiDEs. Within transition in the sensory/ CC related STRiDE was significantly decreased for the reasoning problem solving score of the HC groups. Overall, results indicate that the brain networks tend to be more segregated and less integration in SZ groups.

## Discussion

This study introduces a novel approach to investigating brain connectivity by integrating structural constraints into a data-driven ICA model. The proposed model enables a more nuanced understanding of dynamic functional interactions and their alterations in schizophrenia. Findings highlight the efficacy of this multimodal model in identifying distinct, structurally informed dynamic functional states (STRiDEs) and reveal significant group differences between healthy controls (HC) and individuals with schizophrenia (SZ) across the estimated STRiDEs.

### Methodological Considerations

The connectivity-constrained ICA model marks a substantial improvement over traditional single-modality models by integrating structural priors to yield a more biologically grounded decomposition of functional connectivity. This model addresses the inherent ambiguity in interpreting functional correlations by anchoring them to anatomical constraints, thereby enhancing both the interpretability and robustness of the resulting dynamic functional connectivity states. As highlighted in the Results, the strong correspondence between the estimated STRiDEs and the structural connectivity bases demonstrates the model’s effectiveness in leveraging structural information. This alignment supports a more coherent understanding of brain dynamics, consistent with the view that resting-state functional connectivity is significantly shaped by the underlying structural network, as emphasized by [7].

In addition, multi-modal model evaluation compared to single-modal under the simulation pipeline, highlights a key advantage of the proposed approach in handling noisy data. The estimated STRiDEs consistently identified a higher number of significant connections across varying noise levels, with reduced variability compared to the single-modal method. This suggests that integrating structural and functional information not only enhances the sensitivity of the model but also improves its robustness to noise-related perturbations. In contrast, the single-modal approach demonstrated both lower detection power and greater variability, underscoring its vulnerability in less optimal signal conditions. These findings support the utility of STRiDE in real-world scenarios where data quality may vary and further emphasize the benefits of a multimodal integration strategy for detecting subtle group-level differences.

### Interpretation of Multimodal Brain States

The identification of six distinct STRiDEs reveals a hierarchical organization of brain dynamics, spanning from localized unimodal segregation (e.g., STRiDE1: sensorimotor/visual) to distributed trans-modal integration (e.g., STRiDE3: default mode–cerebellar; STRiDE6: visual–cognitive control). This spectrum reflects established principles of cortical organization, where functional states range from intra-domain coherence to cross-domain interactions [43]. Notably, the alignment of these dynamic states with known structural gradients underscores the influence of anatomical architecture on functional connectivity. The predominance of STRiDE1 suggests a stable, baseline (backbone) configuration supporting essential functions, while less frequent states involving trans-modal integration may support higher-order, context-specific processing [47]. These dynamic transitions likely contribute to cognitive flexibility and adaptive behavior.

### Group Differences in Schizophrenia

Group comparisons between healthy controls and individuals with schizophrenia revealed state-specific dysconnectivity patterns, emphasizing the heterogeneity of functional disruption in schizophrenia. Widespread alterations in sensory and cognitive control domains (STRiDEs1 and 4) and more focal disruptions in DMN and subcortical domains (STRiDEs3 and 6) suggest that schizophrenia impacts distinct functional systems in a dynamic, state-dependent manner. These findings are consistent with prior works demonstrating transient dysconnectivity in schizophrenia [34, 44] and support the view of schizophrenia as a disorder of impaired integration across brain networks [48]. Unlike traditional single-modality studies that often highlight higher-order dysfunction, our structurally constrained approach reveals additional disruptions in sensory and subcortical domains, pointing to early-stage processing deficits. The STRiDE states thus provide a more nuanced view of disease-related alterations by linking dynamic functional abnormalities to structural vulnerabilities.

### Cognitive and Symptom Correlates

The relationship between STRiDEs and cognitive scores underscores the functional relevance of these dynamic states, with group differences linked to attention vigilance, working memory, visual learning, and overall cognitive performance. State-specific associations—such as visual learning with STRiDE2 and working memory with STRiDE6—highlight the role of distinct dynamic connectivity patterns in supporting cognitive functions. These findings align with the view that cognition arises from distributed network interactions rather than isolated regions [49]. Additionally, the identification of STRiDE3 as a symptom-relevant state, showing connectivity changes associated with positive symptoms in schizophrenia, emphasizes the clinical utility of the model. This supports theories linking transient disruptions in front striatal connectivity to psychotic symptoms, consistent with the dopamine hypothesis of schizophrenia [50].

### Temporal Dynamics of Multimodal States

The temporal dynamics of STRiDEs reveal a hierarchical organization of brain activity, with STRiDE1 consistently emerging as the most dominant state—characterized by the highest frequency, dwell time, and occupancy—suggesting a stable baseline configuration from which other, more transient states emerge. This aligns with the concept of a default dynamic state supporting core brain functions, as described by [8]. Notably, strong negative correlations between TC5 and TC2/TC3, along with spatial antagonism in their connectivity profiles, point to a structured interplay between spatial and temporal features of brain organization. STRiDEs with similar or complementary spatial patterns show greater temporal coherence, reinforcing the idea that structural architecture constrains the brain’s dynamic functional repertoire. Differences in dwell time and occupancy between schizophrenia and healthy control groups, along with their cognitive associations, underscore the clinical importance of these temporal features. Disruptions in these dynamics in schizophrenia may reflect impaired flexibility in transitioning between functional states—an essential component of adaptive cognition.

### Speed of Processing

The influence of processing speed on the dynamic interplay among multimodal brain states—particularly the positive association between TC1 and TC6 and the negative association between TC2 and TC4—suggests that efficient coordination between specific networks, especially those involving visual processing (TC6), supports faster cognitive performance. This is consistent with findings by [46], who reported a link between visual attention and processing speed, as well as improvements following visual training. These results underscore the importance of functional integration within and across networks for supporting cognitive efficiency.

### Limitations

While the model effectively captures dynamic functional connectivity, as indicated by strong adjusted R-squared values, some variance remains unexplained. This suggests that additional factors may contribute to brain dynamics. Future work could enhance model performance by incorporating other modalities, such as electrophysiological data, or by applying more sophisticated approaches like dynamic causal modeling (DCM) to explore directional interactions.

## Conclusion

This study highlights the utility of a connectivity-constrained multimodal framework in revealing dynamic functional connectivity patterns and their alterations in schizophrenia. The identification of structurally informed brain states, state-specific dysconnectivity, and cognitive and symptom associations offers a more comprehensive understanding of the disorder. These findings support the integration of structural and functional data in studying brain dysfunction and suggest that future longitudinal and multimodal studies may further inform personalized intervention strategies.

